# Detection of a sequence feature for recursive splicing

**DOI:** 10.64898/2026.04.13.717821

**Authors:** Bixuan Wang, Kevin Yang, Yoseph Barash, Peter S. Choi, Stephen M. Mount, Daniel R. Larson

## Abstract

RNA splicing is directed by *cis*-acting sequence signals interacting with *trans*-acting factors to remove introns from newly transcribed pre-mRNA, joining exons to generate mature mRNA. Splicing happens far more often than observed exon-exon junctions in mRNA. As one contributing process, the spliceosome progressively removes large introns in small segments by ‘recursive splicing’, instead of splicing the whole intron at one time. However, the *cis*-acting sequences associated with recursive splicing have not been identified. Using probabilistic mixture models, we found that recursive splicing occurs more frequently in first introns, which are typically longer, and exhibit a distinct CG-rich sequence feature in the sequences flanking the upstream 5’SS, and depletion of CGs in the downstream polypyrimidine tract. Remarkably, recursive splicing is also more frequent in downstream introns of genes containing first introns with these properties. Mechanistically, these data suggest that early events in RNA synthesis and processing influence the prevalence of recursive splicing for the rest of the transcript. Finally, we developed a sequence-dependent classifier for recursive splicing, which we tested with a novel medium-throughput primer extension assay. In summary, the usage of recursive splicing sites is established at the beginning of RNA synthesis through newly-identified sequence motifs flanking both ends of the first intron.

## Introduction

RNA splicing is a critical regulatory step in eukaryotic gene expression that removes introns from pre-mRNA sequences and ligates exons together. This process allows cells to dynamically define the precise information to be translated into proteins (Deutsch & Long, 1999). The essential mechanism of RNA splicing is carried out by the spliceosome, a large RNA-protein complex, by recognizing various RNA sequence signals (Wilkinson et al., 2020; Will & Lührmann, 2011). Disease-associated mutations can alter splicing by disrupting the conserved splicing site signals or altering the signal recognition components of the splicing machinery (Seiler et al., 2018; G. S. Wang & Cooper, 2007).

Although the conserved steps of splicing and consensus sequences of the spliceosome assembly signals, such as the splicing sites (SS), the branch site, and the polypyrimidine tract, are well characterized, the motifs are highly degenerate and occur much more frequently than real SS (Burge et al., 1999; Will & Lührmann, 2011). Additionally, alternative splicing enabled by the non-continuous exon-intron structure brings both complexity to the transcriptome and diversity to the proteome (Baralle & Giudice, 2017). Typically, splicing regulation is mediated by RNA-binding proteins (RBPs) that recognize various cis-acting elements, and evidence suggests that the same proteins can play opposite regulatory roles when binding at different locations, expanding the landscape of splicing signals. Recent studies have also uncovered co-regulation of splicing and transcription, showing that transcription factors and chromatin structures may impact interactions between trans-acting splicing regulators and pre-mRNA sequences (Begg et al., 2020; Brooks et al., 2015; Fu & Ares, 2014). Due to these complex interwoven regulatory mechanisms, recognizing conventional splicing signals and accurately predicting splicing outcomes remain difficult.

Splicing of long human introns is uniquely challenging. While a combination of strong splicing signals can help with precise and efficient splicing regulation on introns less than 10k nt, this conventional splicing mechanism is less applicable to very large introns greater than 50kb (Gehring & Roignant, 2021; Wan et al., 2021). Potential reasons include the longer transcription time for large introns (Carmo-Fonseca & Kirchhausen, 2014; Coulon et al., 2014) and the reduced efficiency for splice site recognition and spliceosome assembly across great distances. Ultimately, the transesterification reactions of splicing require the splicing sites to be in close spatial proximity, and long genomic distances between them can pose challenges (Fox-Walsh et al., 2005; Gehring & Roignant, 2021). One solution suggested by recent data is that the 5’ SS may be tethered to the elongating polymerase (Zhang et al., 2025). Another solution is recursive splicing (RS), which is a multi-step form of RNA splicing in which introns are removed from pre-mRNA sequences as multiple smaller pieces marked by recursive splicing sites (Duff et al., 2015; Gehring & Roignant, 2021; Hatton et al., 1998; Pai et al., 2018; Sibley et al., 2015). Intermediate steps in RS include recognition of splicing signals and removal of intron subsequences by the spliceosome, leaving behind the rest of the intron sequences with additional recursive splicing sites to be spliced out subsequently (Gehring & Roignant, 2021; Hatton et al., 1998; Pai et al., 2018). Such a unique splicing process may help maintain splicing fidelity over large distances and add an additional layer of complexity to gene expression regulation. However, the molecular mechanisms involved are not understood, and it is unknown what distinguishes these re-splicing sites from those used in the final exon-exon junction of the mRNA.

Recent studies indicate that recursive splicing occurs extensively in humans. Though initially documented in both Drosophila and mammals, RS was thought to be suppressed by the exon-junction complex (EJC), which marks the completion of splicing in humans (Blazquez et al., 2018; Boehm et al., 2018). However, advanced techniques such as dynamic single-molecule imaging and high-throughput sequencing of nascent RNA and splicing lariats have revealed genome-wide RS site usage in humans (Wan et al., 2021). Single-molecule FISH revealed that the RS process was carried out using multiple RS sites before the downstream canonical 3’SS was transcribed in large introns longer than 10kb. These findings have been confirmed with in-depth analysis of the Sequence Read Archive (Duan et al., 2024). A model that recursive splicing is pervasive and on-pathway is supported by data from lariat sequencing (Wan et al., 2021). Although these RS sites and the resulting intermediates were observed to be utilized stochastically, there is likely sequence encoded information that has yet to be identified. In summary, the cis-acting sequence information and the trans-acting RBPs guiding RS site usage remain unknown. Moreover, although the spliceosome appears to be far more active than previously appreciated, why the spliceosome ultimately stops at the ‘correct’ exon-exon junction is not understood.

Here, we investigate cis-acting and trans-acting RS regulatory elements in nascent RNA-Seq data (Wan et al., 2021). By comparing sequences associated with canonical and recursive splicing sites, we found that RS occurs more frequently in first introns, which are typically longer. Using sequence k-mer enrichment analysis methods from natural language processing, including a mixture model called Latent Dirichlet Allocation (LDA, (Blei et al., 2003; Wang & Mount, 2024)), we found that a CG-rich motif is enriched in the vicinity of the 5’SS. Notably, this same motif was selectively depleted upstream of 3’ SS in first introns of genes which show recursive splicing, constituting a new type of pyrimidine tract. Surprisingly, we also found that genes where the first introns use RS are more likely to have RS in downstream introns. Moreover, whole-genome bisulfite sequencing indicates that the first exons of genes with RS have lower CpG methylation rates. With the sequence features discovered, we trained a random forest classifier to predict recursive splicing events with an accuracy of over 84% in the first introns and over 80% in downstream introns, which we then tested experimentally using a medium throughput primer extension assay. We propose that the use of RS sites throughout a gene is influenced by sequence elements early in the transcription unit that presumably act by recruiting factors which remain associated with polymerase throughout transcription.

## Results

### Identification of sequence and gene architecture features associated with recursive splicing

Our approach to identifying RS signals began with systematic analysis of nascent RNA-Seq data (Wan et al., 2021) compared with mature RNA-Seq of the same cell line (Donovan et al., 2024). We first searched for potential RS junctions from nascent RNA-Seq collected during a pulse-chase experiment after a DRB-washout. This experiment consists of treating cells for 3 hours with the transcription elongation inhibitor DRB, which effectively clears RNA polymerase II from genes. Following washout of the inhibitor, nascent RNA is collected at multiple time points to capture intermediates in transcription and splicing of nascent transcripts. We selected splicing junctions that overlap with annotated human introns and have at least 3 unique supporting reads. Nascent RNA samples show a higher representation of unannotated splice junctions than do mature RNA samples (Figure 1A). Splicing junctions found in the nascent RNA samples are much smaller, on average, than annotated junctions found in mature RNA, and junctions present in both nascent and mature samples are the smallest. (Figure S1A), implying that splicing intermediates are on their way to fully spliced mRNA. This observation is consistent with the stepwise recursive splicing process proposed previously, where large introns are spliced out in pieces (multi-step) rather than as a single lariat (single step) (Gehring & Roignant, 2021; Hatton et al., 1998; Wan et al., 2021). Next, we compared the coordinates between unannotated splicing junctions and annotated introns. About 78% of the unique unannotated splicing junctions are subsequences from annotated introns, and the sizes of the splicing junctions are similar to introns that were spliced out completely in single-step splicing in nascent RNA samples, indicating a preferred intron length of about 1k nt for individual splicing events (Figure S1B, 1B). These observations support the hypothesis that the spliceosome progressively removes introns in small pieces through recursive splicing instead of splicing a whole intron at once (Figure 1C).

**Figure 1.**
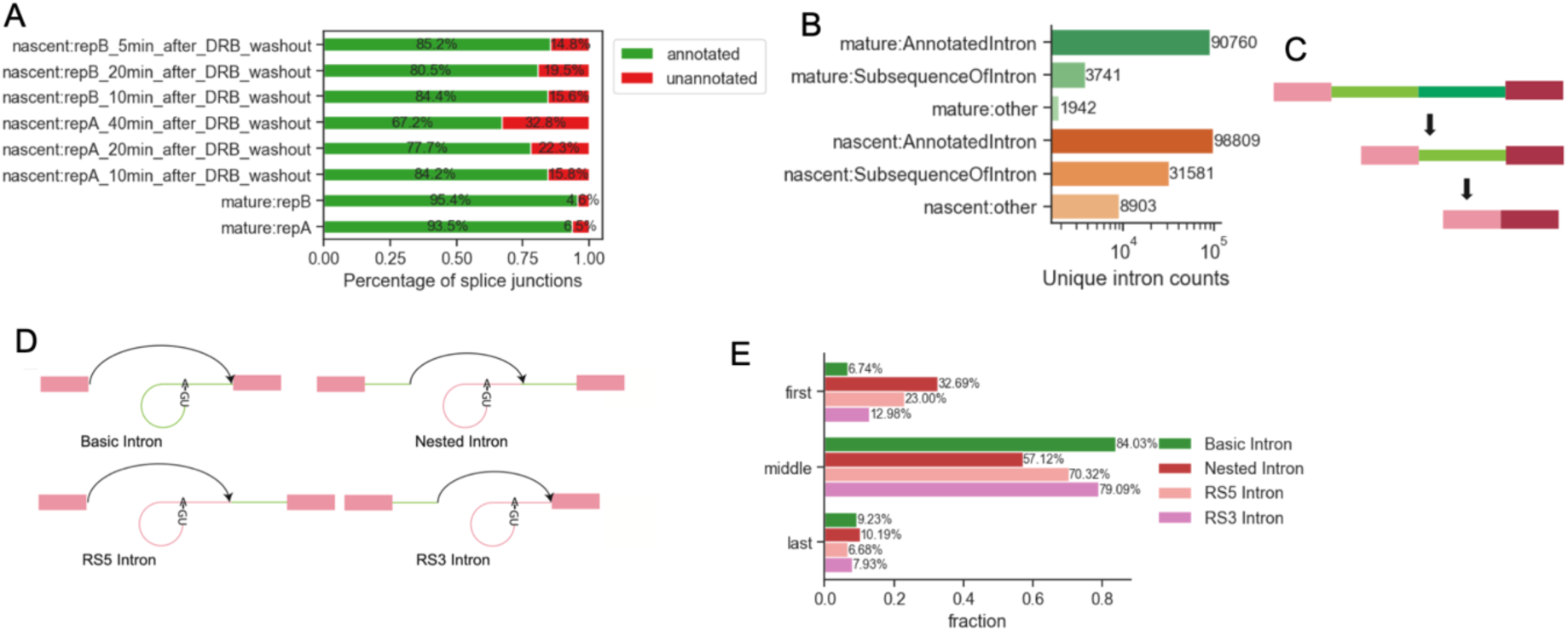
Nascent RNA-Seq revealed 4 types of introns. A. Fractions of annotated and unannotated splicing junctions from nascent and regular RNA-Seq. Fractions are based on annotations of splicing junctions from STAR output with unique spanning reads >=3. B. Number of unique introns in regular and nascent RNA-Seq. Unannotated splice junctions are labeled based on if they are subsequences of annotated introns (subsequence or other). Nascent RNA-Seq shows more unannotated splice junctions where most of them are subsequences of annotated introns. C. Recursive splicing model. The spliceosome progressively removes the intron sequence by splicing the 3’ segment (dark green) in the first step. Recursive splicing stops when the whole intron is removed. D. 4 types of introns identified from nascent RNA-Seq. Basic introns are annotated introns without recursive splicing junctions from STAR output. RS5 introns are introns show splicing junctions with annotated 3’SS and unannotated 5’SS downstream of the annotated 5’SS. RS3 introns are introns with annotated 5’SS and unannotated 3’SS upstream of the annotated 3’SS. Nested introns are introns with pairs of unannotated SS. E. Fractions of four types of introns as the first, middle, or last introns of transcripts. Compared with basic introns, RS introns are more likely to appear as the first introns.

This sequence analysis allowed us to define classes of recursive splicing. From the alignment between unannotated splicing junctions and annotated introns, we summarized 3 types of splicing sites:

1. Canonical splicing sites from human genome annotations.
2. Recursive 5’SS that pairs with canonical 3’SS or recursive 3’SS.
3. Recursive 3’SS that pairs with canonical 5’SS or recursive 5’SS.

Pairing these canonical/recursive 5’ and 3’ splicing sites describes 4 types of introns: basic introns without RS sites, RS5 and RS3 introns with one recursive and one canonical splicing site, and nested introns marked by both recursive splicing sites (Figure 1D). Splicing junctions with single RS sites have similar lengths to basic introns in nascent RNA samples, while nested introns are the smallest and rarest splicing junctions (Figure S1C). We note that RS sites are expected to act as both 5’ and 3’ SS with conserved AGGU motifs. Yet, we found that 56.4% of RS5 Introns and 62.6% of Nested Introns show AGGU/C at recursive 5’SS, and only 25.3% of RS3 Introns and 37.5% of Nested Introns show AGGU/C at the recursive 3’SS (Figure S1D). Positional weight matrices show slightly lower information content in the 3 bases downstream of recursive 5’SS and the polypyrimidine tract of recursive 3’SS in nested introns (Figure S1E-H). Annotation of the intron position of genes indicates that the first introns from multi-exon genes, typically very long, are more likely to have RS than the downstream introns (Figure 1E). In summary, we identify four classes of introns from nascent RNA-Seq with distinct RS site usage and observe that RS is more likely to occur in the first intron of genes.

We hypothesized that RS in first and downstream introns might have different mechanisms and therefore did sequence feature analysis for them separately. K-mer composition modeling was calculated using a probabilistic model called the mixture model (Blei et al., 2003; Pritchard et al., 2000; Wang & Mount, 2024), which represents each sample -- such as a sequence -- as a mixture of complex features, which are mixtures of simple components, such as k-mers. Click or tap here to enter text.We modeled sequences at 5’SS and 3’SS separately using sequences flanking the annotated splicing sites (Figure 2A). In addition to canonical splicing sites and recursive splicing sites, we searched for random AGGU motifs in related introns as pseudo-RS sites, which helped us to distinguish the features of RS sites besides the tetramer motif (Figure S2A-S2H). We aligned sequences from 30bp upstream to 50bp downstream of 5’SS and upstream 50 bp to downstream 30 bp of 3’SS at the splicing sites and counted the tetramers at each position with a sliding window of 6 bp (Figure 2A, methods). Based on previous experience (Wang & Mount, 2024) we aimed to find six features or ‘topics’ using mixture models for the first intron 5’SS, first intron 3’SS, downstream intron 5’SS, and downstream intron 3’SS separately. The result describes the k-mer composition of annotated SS at each position as a mixture of six features; each feature is a mixture of k-mers with distinct distributions (Figure 2A, methods). In all sequence types, mixture models discover the strong signal of AGGU as expected, which represents potential recursive splice sites shown as a topic enriched between −8bp upstream to the splice site position as we used a sliding window of 6bp to smooth the signals (Topic 3, light green, Figure 2B-2E, S2A-S2H).

**Figure 2.**
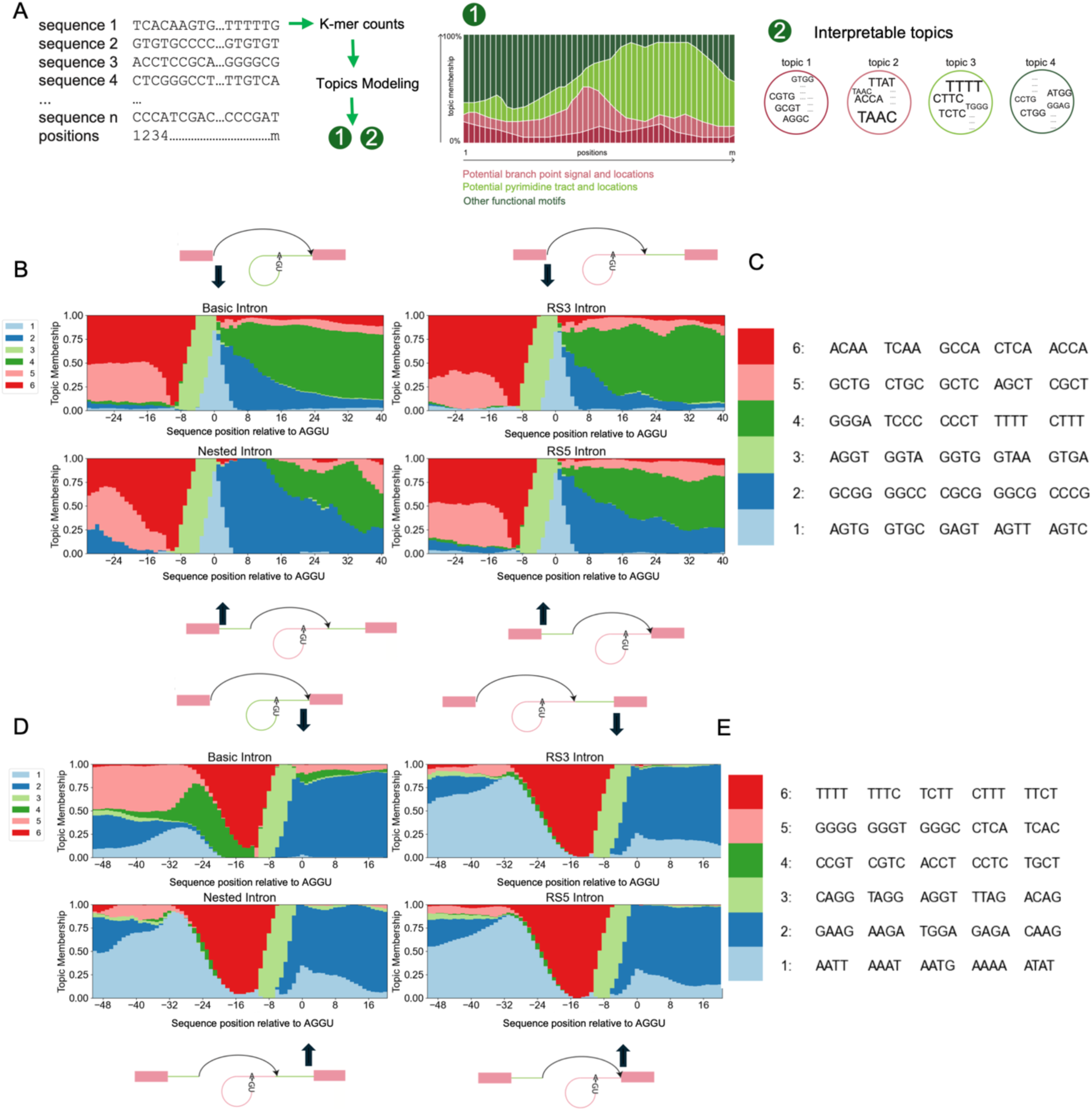
Mixture models uncovered potential cis-acting factors at canonical 5’SS. A. Evaluation of potential cis-acting recursive splicing factors by mixture models. A mixture model takes k-mer counts of each sequence sample as input and returns topic membership of the sequence samples and a list of interpretable topics. Each topic is a list of scores corresponds to all possible k-mers. Sequence topic memberships help reveals sequence function represented by enriched motifs. B. Structure plot of feature/topic distribution of regions from 30bp upstream to 50bp downstream of canonical 5’SS (sliding widow=6bp) in the first introns. Mixture model is fitted with k-mer counts at different positions (x-axis) across sequences. The y-axis measures the memberships of topics. The position 0 is the 5’SS, which is enriched (due to the sliding window, from −8∼0) in topic 3, where the top 1 k-mer is AGGU. C. Top ranked k-mers in the 5’SS model in B. D. Structure plot of feature/cluster distribution of regions from 50bp upstream to 30bp downstream of canonical 3’SS (sliding widow=6bp) in the first introns. The position 0 is the 3’SS which is enriched in topic 3 of AGs. E. Top ranked k-mers in the 5’SS model in D.

Grouping first introns by whether RS was observed helped us identify an RS-related motif at the canonical 5’SS. In the region flanking the canonical 5’SS in the first introns, higher CG-rich feature enrichment (Topic 2, dark blue, Figure 2B, 2C, S2A, S2B) is observed in the sequences with recursive splicing sites downstream, especially in Nested Introns and RS5 Introns, while in the region of other canonical splicing sites, such signal is much weaker in a grade of membership plot. Comparison of recursive and canonical 5’SS of the first introns failed to reveal any striking differences beyond the trivial difference between exon-intron boundary and deep intron sequence contexts. Such differences are expected based on our definition of annotated splice sites (Figure S2A, S2B). We then compared recursive 5’SS and random AGGU and observed an AT-rich feature (Topic1, dark red, Figure S2A, S2B) in flanking sequences, consistent with the fact that both types of sequences are in deep intron regions. However, this intronic feature is more enriched in the downstream sequences of recursive 5’SS than in the upstream sequences. Another difference between recursive 5’SS and random AGGU is the light enrichment of a GC- and AG-rich feature (Topic 5, light red, Figure S2A, S2B) at the upstream of the SS/AGGU location, which is a feature also observed upstream of annotated SS in all four types of introns. These observations suggest that the sequences flanking a recursive 5’SS have a context resembles exon-intron boundaries.

Comparison of recursive and canonical 3’SS of first introns also reveals differences. Both sites share the same features upstream of the 3’SS, but downstream of the canonical 3’SS there is enrichment of an AG feature (Topic 2, dark blue, Figure S2C, S2D) while downstream of recursive 3’SS is more enriched in a feature of GCT/TCG (Topic4, dark green, Figure S2C, S2D). Again, this difference could result from sequence context differences between exons downstream of the canonical 3’SS and the deep intron sequences. Yet, from sequences flanking the annotated 3’SS of the first introns, we observed a G-rich pattern located upstream of the 3’SS that shows a stronger signal in the introns with no evidence of recursive SS (Topic 5, light pink, Figure S2C, S2D). At the upstream of the canonical 3’SS of RS introns, the fraction of such a feature is much lower, while another AT-rich feature is stronger in both upstream and downstream of the canonical 3’SS (Topic 1, light blue, Figure S2C, S2D). To take a closer look at the feature of annotated 3’SS of the first introns, we fitted another mixture model using only sequences flanking the annotated 3’SS. The new model revealed that besides a G-rich pattern that we previously observed (Topic 5, light pink, Figure 2D, 2E S2C, S2D), the introns with no RS evidence show a smaller fraction of a feature representing strong polypyrimidine tract (Topic 6, dark red, Figure 2D, 2E) but have a unique feature of CTG/CTC (Topic 4, dark green, Figure 2D, 2E), which indicates a different polypyrimidine tracts present upstream of the canonical 3’SS in RS introns and canonical introns. In contrast, no significant differences were observed in downstream introns between recursive and basic introns (Figure S2E-S2H). Counter intuitively, the RS introns appear to be associated with longer and stronger polypyrimidine tracts than basic introns, where purines tend to be present starting at ∼ 20 nt upstream of the 3’ SS.

Our data suggested the existence of two features associated with RS events: 1) a CG-rich feature flanking the beginning of the first introns and 2) a different pyrimidine tract upstream of the 3’SS of the introns. To further probe this signal, we looked at meta-gene alignment of the first and downstream introns. Meta-gene profiling on the upstream exon, the intron, and the downstream exon sequences indicates that the beginning of the first intron with recursive 5’SS has relatively higher GC content than the corresponding region of the first introns without recursive 5’SS (Figure 3A, 3B). In contrast, the remaining regions of the first introns and all three areas of downstream introns indicate the opposite: RS-related sequences have lower GC content (Figure 3A, 3C). A complementary metric -- tetramer enrichment evaluation -- is likewise consistent with this observation (Figure 3D, 3E).

**Figure 3.**
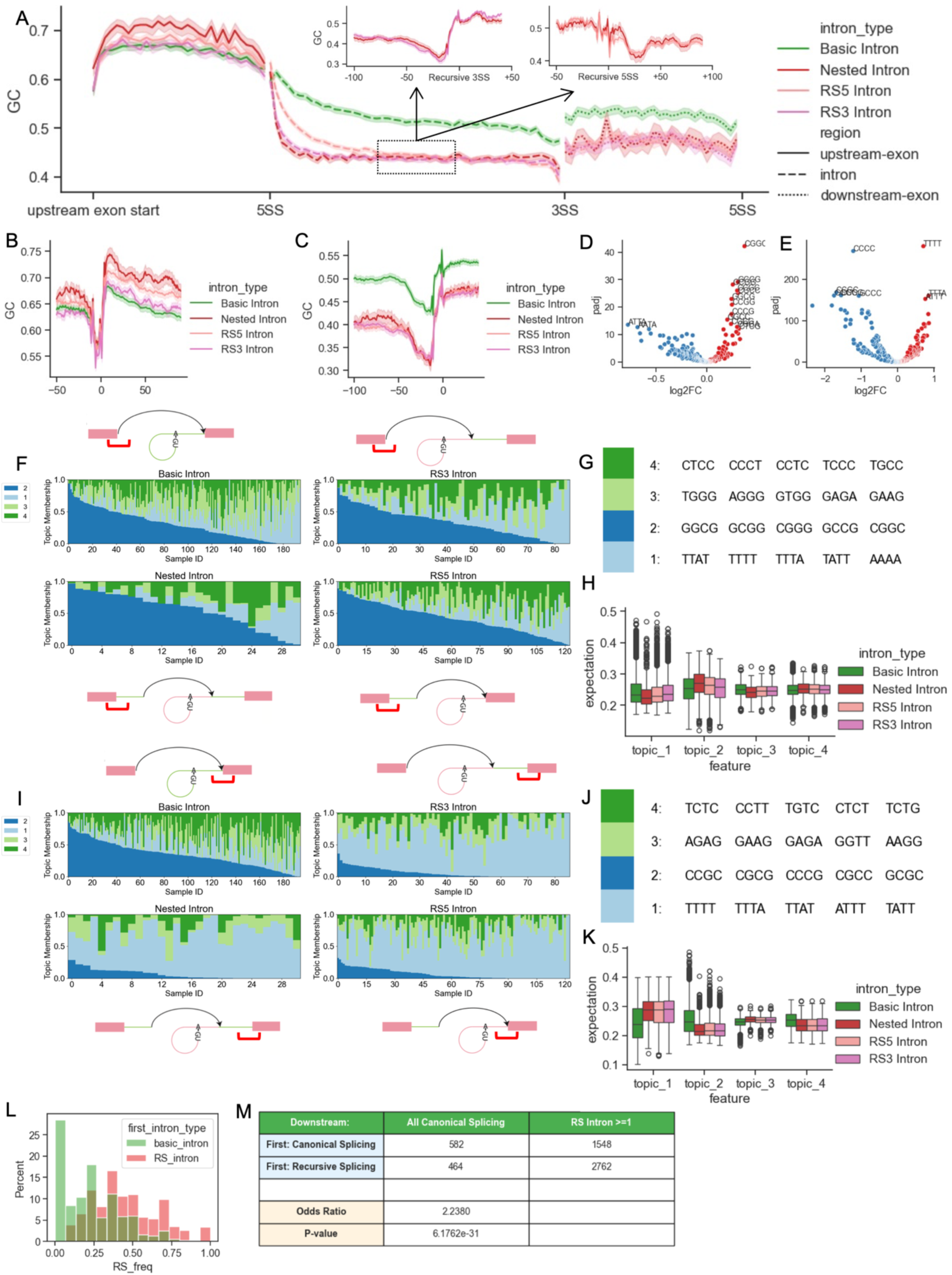
Sequences of the annotated 5’SS and 3’SS of the first intron indicate recursive splicing site usage. A. Meta-gene plot of GC content of 4 types of first introns and adjacent exons. RS introns have relatively lower GC content in the intron and downstream exon sequences. Nested introns show higher GC content in the upstream exon sequences. Smaller windows are focusing on recursive SS, where lower GC contents are shown upstream of recursive 3’SS and downstream of recursive 5’SS. B. GC content in the region from upstream 50bp to downstream 100bp of annotated 5’SS in the first introns. C. GC content in the region from upstream 100bp to downstream 50bp of annotated 3’SS in the first introns. D. Volcano plot of tetramer enrichment in the sequences in region shown in B. Multi-test correction is applied to the log2FC calculation of k-mer enrichment. Red dots enriched in CGs are tetramers enriched in introns with recursive splicing sites. E. Volcano plot of tetramer enrichment in the sequences in region shown in C. Multi-test correction is applied to the log2FC calculation of k-mer enrichment. Red dots enriched in Ts are tetramers enriched in introns with recursive splicing sites. F. Structure plot of distribution of features in the region from 50bp upstream to 50bp downstream of annotated 5’SS in the first introns. Mixture model is fitted with k-mer counts of sequence samples (x-axis, Sample ID), and each sample contains 15 sequences. The y-axis shows memberships of each topic. G. Top k-mers in the model in F. H. Distribution of single sequence topic expectation values calculated using model F. Nested introns and RS5 introns are slightly more enriched in topic 2 features. I. Structure plot of distribution of features in the region from 50bp upstream to 50bp downstream of annotated 3’SS in the first introns. J. Top k-mers in the model in I. K. Distribution of single sequence topic expectation values calculated using model I. Basic introns show more enrichment in topic 1 and less enrichment in topic 2 than all 3 types of RS introns. L. Distribution of the fraction recursive splicing introns in all the expressed downstream introns. Colors indicate if the first introns in the transcripts show recursively splicing junctions in nascent RNA-Seq. M. Contingency table and Fisher’s Exact result of evaluating the effect of first intron splicing on downstream introns.

To further explore potential motifs in sequences flanking canonical 5’SS and 3’SS in the first introns and to confirm the GC content comparison is not biased by small subsets of GC-rich sequences, we applied mixture models on k-mer compositions. The sequence samples are composed of 30bp in the exon region and 50bp in the intron region (e.g. 30bp upstream to 50bp downstream of annotated 5’SS), each of which is a group of 15 sequences randomly selected from the same intron types (Figure 3F-3K). As a generative model, the mixture model presents the feature interpretation matrix as the probabilities of observing each k-mer in each feature. To evaluate the mixture feature/topic distribution in every single sequence, we calculated the expectations of observing each feature in those sequences (Figure 3H, 3K). The topic membership distribution shows higher fractions of topic 2, in which CG-rich k-mers are distinctively expressed compared with other features, in the sequences flanking the annotated 5’SS upstream of potential recursive 5’SS from either RS5 or Nested introns (Figure 3F-3H). In contrast, the mixture model fitted with 3’SS sequences revealed that the first introns that have RS show a lower CG-rich feature, which is also represented as topic 2, than canonical introns (Figure 3I-3K). Though the major difference at the canonical SS between intron with and without RS lies in the AT-rich and CG-rich feature distributions, we find that mixture modeling also assigned higher memberships of topic 4, a CCT/CCA-rich feature, to RS-related sequences at the 3’SS (Figure 3I-3K), suggesting that the signal at the canonical SS may not be explained by GC content alone.

Although we identified the CG-rich features associated with RS in the first introns by multiple approaches, this feature was not readily observed in downstream introns where RS was present. In fact, the differences in the topic distribution in downstream introns are more subtle and difficult to interpret (Figure S2E-S2H). However, we noticed through gene annotation of the splicing junctions that more downstream introns are recursively spliced when RS events are observed in the corresponding *first* introns (Figure 3L). We therefore hypothesized that recursive splicing in the first intron makes downstream introns more likely to be recursively spliced. We analyzed transcription units with at least three introns detected in the RNA-Seq data using a Fisher’s Exact test and observed a significant association between RS in first and downstream introns with an odds ratio of 2.2. Moreover, we noticed that the number of transcription units without recursive splicing is much smaller than transcription units with at least one recursively spliced intron (Figure 3M).

The coupling between first intron RS and downstream intron RS suggests that the sequence features near the first canonical 5’SS (the CG-rich feature) and 3’SS (difference in the polypyrimidine tract) could propagate far into the gene. To test for this effect, we repeated the mixture modeling by classifying the first introns into 6 types of transcripts: in addition to 4 of them based on the contingency table, we gathered the first introns where the splicing status is unclear from nascent RNA-Seq *but* RS events in downstream introns are known (Figure S3A). Mixture feature distribution showed statistically insignificant, slightly higher memberships of CG-rich feature in sequences flanking the first canonical 5’SS from transcripts showing RS in the first intron than transcripts with same splicing status downstream but no RS in the first intron (Figure S3B-S3D). However, the same comparison at the canonical 3’SS showed that when the first intron splice status is fixed, the transcripts with downstream RS or no RS have significantly different mixture feature distributions, including the transcript units where the first intron splice status is unknown (Figure S3E-S3G). In summary, sequence features in the first introns of genes are associated with the splicing activity for downstream introns.

One prototypical functional CG-rich motif is the CpG island, a cluster of CG methylation sites that is frequently found at the 5’ end of genes, including first exons (Anastasiadi et al., 2018). To assess whether the CG sites associated with this signal are subject to methylation, we calculated the average CpG methylation rate of each exon using the Whole-Genome Bisulfite Sequencing profile of the A549 cell line from ENCODE ((Dunham et al., 2012; Luo et al., 2020); Figure 4A). The methylation rate distribution indicates that at the first introns, the exons upstream of RS introns (the first exons of the genes) are less methylated than exons upstream of the introns with no RS events (Figure 4B). This effect is evident across all classes of RS junctions (nested, RS3, RS5). Moreover, we classified the first introns based on whether RS is observed in the downstream introns of the same genes. The first exons from genes that have RS events in the downstream introns showed less CpG methylation than genes that show no RS evidence (Figure 4C). Overall, we show that the CG-rich motif with a low CpG methylation rate adjacent to the first canonical 5’SS is associated with RS of the genes. This intriguing association between the CG-rich motif, low DNA methylation, and splicing outcomes may suggest selection pressure to maintain the motif (see discussion).

**Figure 4.**
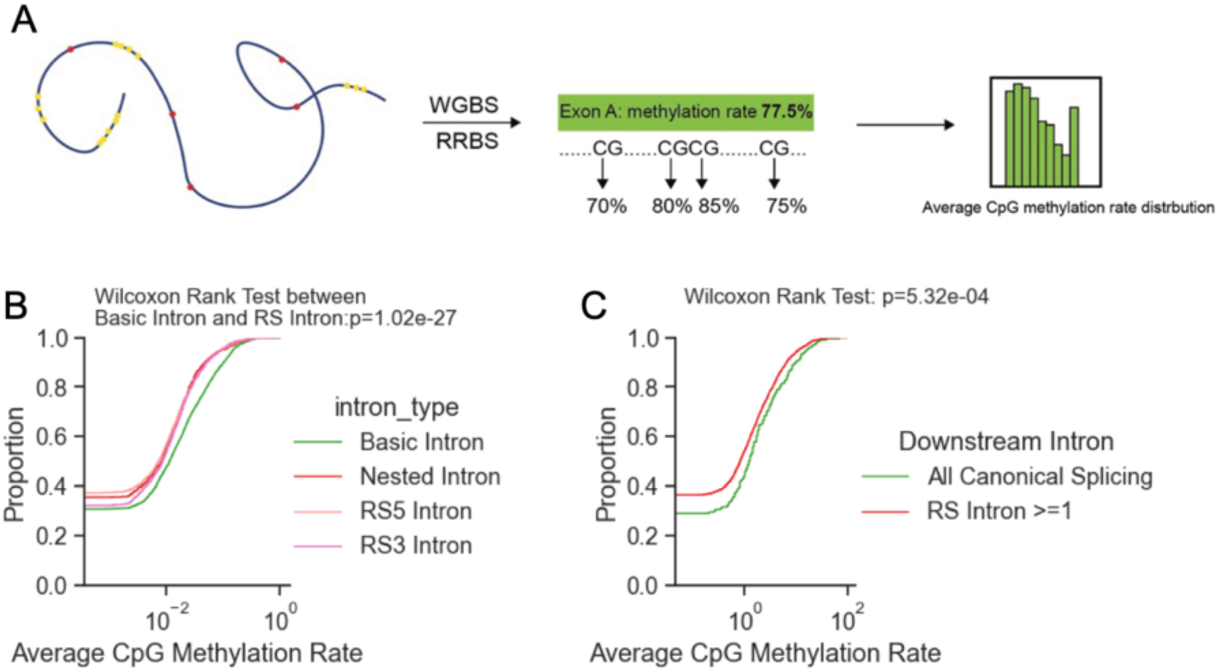
Recursive splicing is associated with low CpG methylation in the first exons. A. Diagram of the calculation of exon average CpG methylation rate. CpG methylation profile of A549 cell line from ENCODE is used to calculate average CpG methylation rate of each exon. B. Cumulative distribution of upstream exon methylation rate for 4 types of introns as first introns of genes. Wilcoxon rank test is applied for significance test. C. Cumulative distribution of first exon methylation rate for transcripts with or without recursive splicing in downstream introns.

In summary, we identified sequence signatures of recursive splicing. These sequence signatures were discovered through mixture models and latent Dirichlet allocation and are: 1) a CG-rich motif which is present near the 5’SS of first introns, and 2) a different, significantly stronger polypyrimidine tract of the first introns. These sequence signatures not only appear to predict RS in first introns but also the probability of RS in downstream introns. These observations suggest it might be possible to quantitatively predict RS and potentially gain mechanistic insight.

### Classifier for accurate recursive splicing prediction

In order to develop and test a classifier for RS prediction, we used the potential cis-acting elements to train random forest classifiers to predict RS events in first and downstream introns separately. For the first intron RS prediction, we selected 4 regions for mixture modeling: the upstream exons, 50 bp sequences downstream of the canonical 5’SS, 50 bp sequences upstream of the canonical 3’SS, and the downstream exons (Figure 5A). We found that a combination of mixture feature expectations, intron sizes, and upstream exon CpG methylation information could correctly classify 84% of the test sequences based on the Area Under the Curve (AUC) from the Receiver Operating Characteristic (ROC) Curve (Figure 5B). Topics of sequences flanking the canonical 3’SS are among the top important features of the classifier, which is consistent with the mixture modeling in our previous results (Figure 5C). Moreover, our model confirmed the findings from previous studies that recursive splicing occurs more often in longer introns and in highly expressed genes, and provided an additional quantitative estimate of the extent to which these factors affect the usage of recursive splicing sites. We also note that ‘intron length’ is an emergent property which may itself contain contributions from the other features. By selecting matching subsets from basic introns and RS introns (Figure S4A), we confirmed that among introns of similar lengths and TPM, features of first intron sequences help predict RS with an accuracy of 75% (Figure S4B).

**Figure 5.**
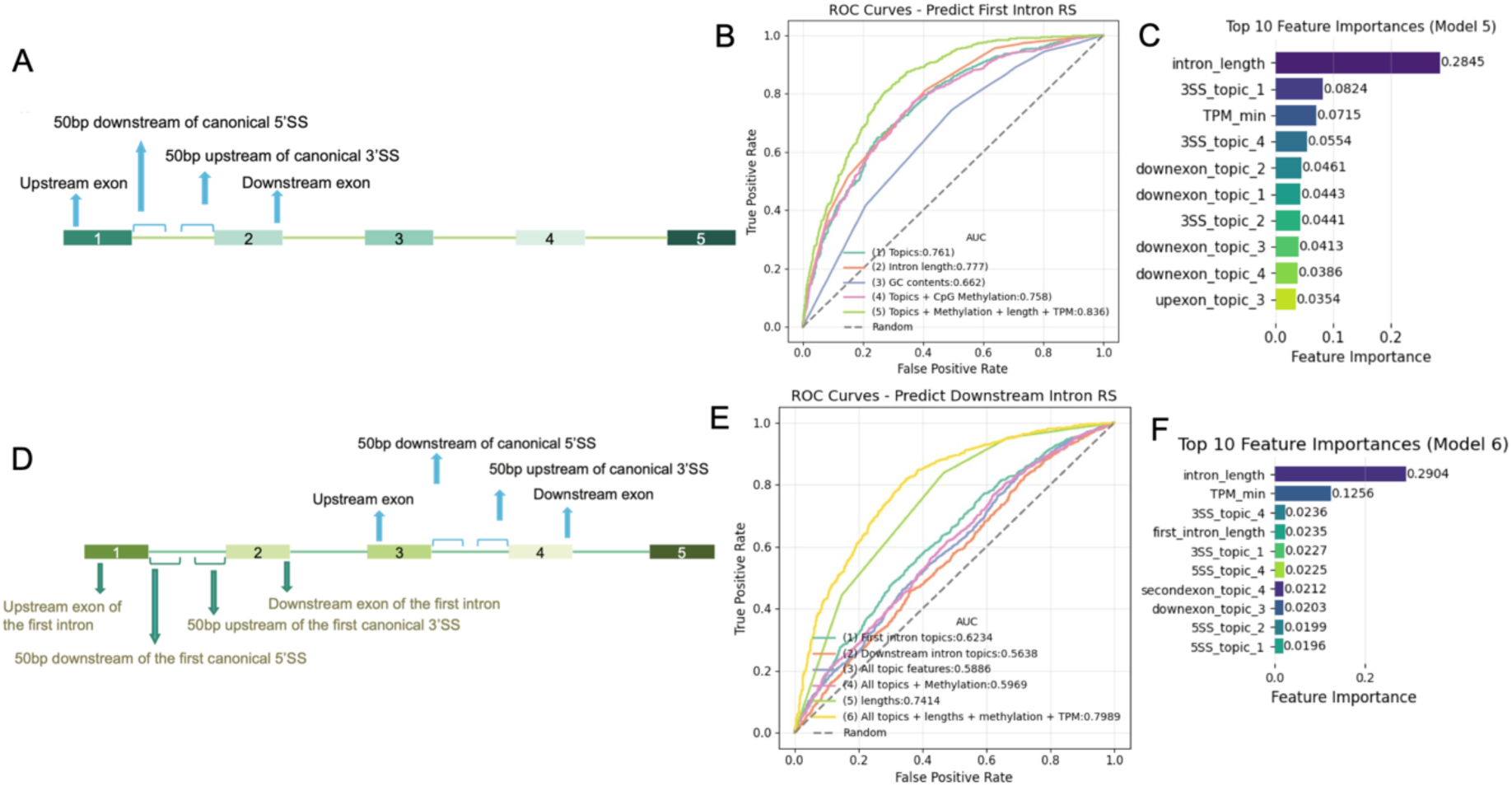
Computational modeling of sequence features aids in the accurate prediction of recursive splicing. A. Sequences selected for mixture modeling and random forest classifier fitting to predict recursive splicing in the first introns. Sequence topic/feature memberships from 4 mixture models fitted with 4 groups of sequences are used for recursive splicing predictions. B. ROC curve of predicting recursive splicing of first introns. Curves of colors are showing performances of random forest classifiers using different features. C. Top 10 features and importance values ranked by random forest classifier (green curve in B). D. Sequences selected for mixture modeling and random forest classifier fitting to predict recursive splicing in the downstream introns. E. ROC curve of predicting recursive splicing of downstream introns. F. Top 10 features and importance values ranked by random forest classifier (yellow curve in E).

To predict RS events in downstream introns, we prepared the features of the corresponding first introns of the same genes in addition to the features of the downstream introns (Figure 5D). Performance evaluation of downstream RS prediction indicates that the classifier can achieve an AUC of 80% (Figure 5E). Similar to the first intron RS prediction, the performance relies heavily on the intron length and gene expression level, which have combined importance score over 40%, followed by the length of the first intron as the fourth important feature, which is the most important feature to predict first intron splice status (Figure 5F). To avoid the impact of those features in RS prediction, we picked matching sets similarly as we did for first introns and confirmed the prediction accuracy using sequence features is about 61% (Figure S4C, S4D), driven by the feature of the first introns, consistent with our finding that the usage of recursive splicing sites in the downstream introns is associated with splicing status of first introns of the genes. In summary, the potential cis-acting features are able to predict the presence of RS in the genes expressed in nascent RNA-Seq data with an accuracy of over 85% for first introns and 80% for downstream introns.

We next sought to generate a new dataset as a test of the classifiers. Local splicing variation sequencing (LSV-seq) is a medium-throughput method targeting splicing variations upstream of selected exons with optimized primers (Yang et al., 2025). In this context, the goal was to identify groups of introns, computationally design stringent primers, carry out reverse transcriptase primer extension assays, and sequence the resulting cDNA. We hypothesized that some introns are removed through recursive splicing which are not captured by RNA-Seq due to fast splicing reactions or low expression, but which can be detected by LSV-seq. Furthermore, we attempted to query a range of expression levels, intron lengths, and positions within transcripts. We selected four groups of introns for targeted recursive splicing validation by LSV-seq: (1) introns that show recursive splicing junctions from RNA-seq and have corresponding high scores from the classifier (n=91), (2) introns that show zero recursive splicing junctions from RNA-seq and have corresponding low scores from the classifier (n=86), (3) introns that show zero recursive splicing junctions but have high probabilities of recursive splicing calculated by the classifier (n=39), and (4) introns that were not expressed in nascent RNA-Seq but have high probabilities of recursive splicing (n=11) (Figure 6A, B).

**Figure 6.**
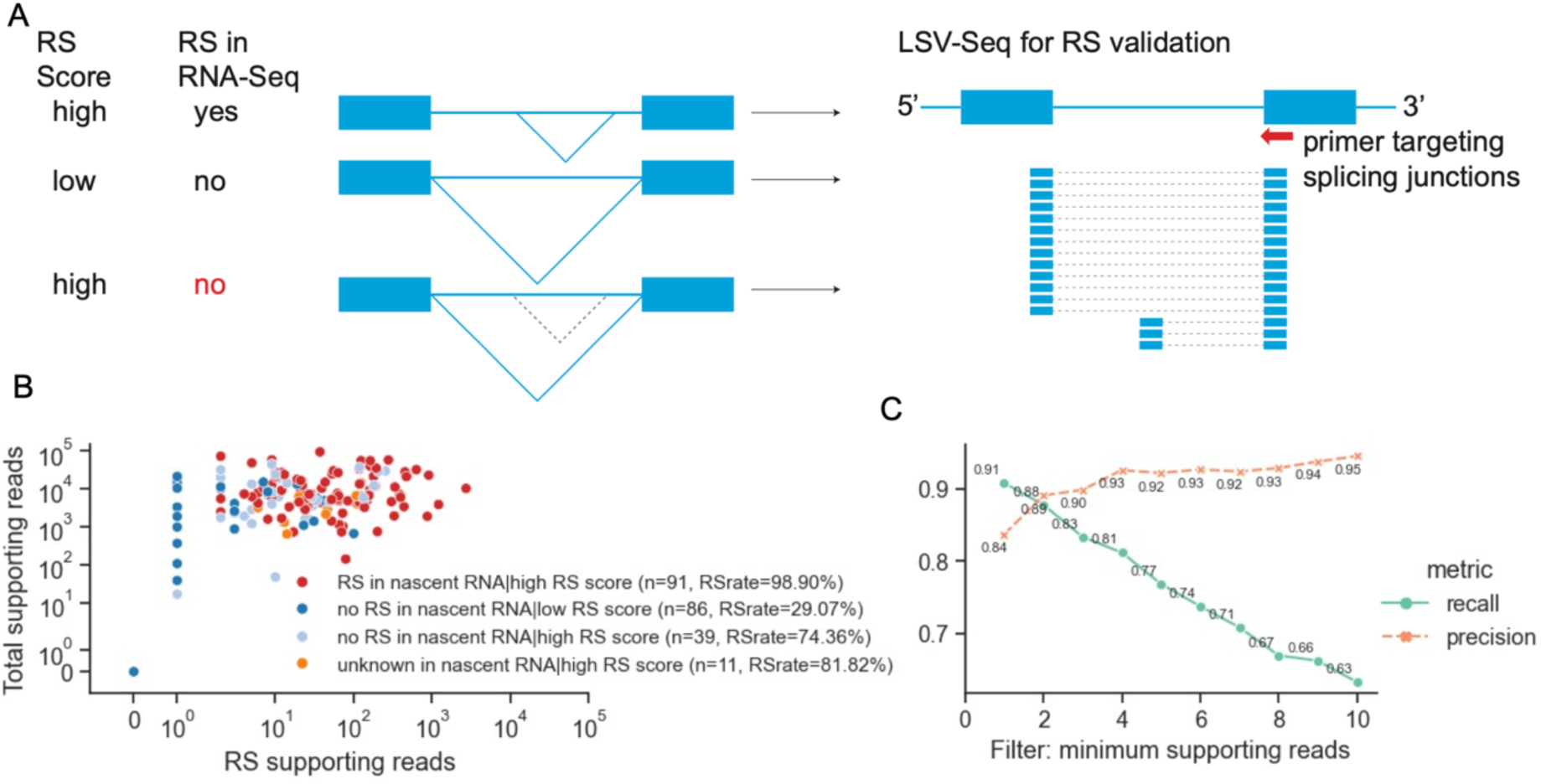
Validation of recursive splicing prediction by LSV-Seq. A. Diagram of LSV-Seq validation of recursive splicing from 3 groups of introns: (1) both random forest classifier and nascent RNA-Seq data indicate RS in the intron; (2) both random forest classifier and nascent RNA-Seq data indicate no RS; (3) the random forest classifier predicts RS in the intron while nascent RNA-Seq show no RS or low gene expression. LSV-Seq is applied to search for potential recursive 5’SS usage. B. Scatterplot of the read counts showing canonical and recursive splicing in 4 groups of introns. Unique supporting reads for recursive splicing and the intron region are shown. C. Precision-recall lineplot of the recursive splicing validation accuracy by using different read count filtering. Precision and recall are calculated using random forest classifier output as truth and LSV-Seq output as observation, using introns with supporting reads>=10.

The classifier does exceptionally well at predicting previously undetected RS events. Specifically, LSV-seq reveals potential recursive splicing junctions in 98.9% of introns in group (1) (90/91), 74.36% of introns in group (3) (29/39) and 81.82% (9/11) of introns in group (4). Introns in group (2) are predicted to have low probability of recursive splicing, and we detect 29.07% (25/ 86) (Figure 6B). Notably, the success rates in test groups (3) and (4) suggest that many RS events were missed in previous datasets but are recovered in the targeted LSV-seq approach across a range of expression levels. To more precisely test the classifier, we label each intron as recursively-spliced or not recursively-spliced based on the classifier predictions and measure the performance of the LSV-seq output by filtering the introns with supporting reads (RS read threshold, Figure 6C). The performance metrics show precision over 89% and recall values over 88% when applied to only introns with more than two unique supporting reads. Increasing this threshold 4x has a negligible effect on precision and degrades the recall to ∼ 66%. In conclusion, we developed and tested an RS classifier with an accuracy of 84% in first introns and 80% in downstream introns. Moreover, the features of this classifier have mechanistic predictions for the regulation of RS.

### Predicting candidate trans-acting regulators of regulate recursive splicing

Accurate and efficient intron removal relies on multiple RBPs, which regulate the splicing process by recognizing and binding to specific motifs in the RNA sequence (Brooks et al., 2015; Dominguez et al., 2018a; Van Nostrand et al., 2017, 2020). To identify trans-acting RBPs involved in RS regulation, we used the binding preference inferred by RNA Bind-N-Seq (Dominguez et al., 2018b; Lambert et al., 2014). RBNS is a high-throughput technique characterizing RNA sequences bound by RBPs, which can be used for the comparison of k-mer counts between the RBP and control library to reflect the binding preferences and regulatory functions. Our strategy was to identify candidate RBPs that might interact with either the GC-rich signal or the modified polypyrimidine tract as those with a preference for the driving k-mers identified through mixture models.

To identify potential differential binding activities in various regions associated with recursive splicing, we first summarized a k-mer protein matrix using the R values of RBNS from ENCODE (Dunham et al., 2012; Luo et al., 2020; Van Nostrand et al., 2017, 2020), which are ratios of bound k-mers between protein and control libraries. To map the R-values to RS sequences, we calculated the fractions of k-mers in regions 75 bp upstream to 75 bp downstream by grouping every 15 bp as one unit. We multiplied the position k-mer matrix with the k-mer protein matrix to get a position protein R-value matrix (Figure 7A). To identify proteins with differential binding activities between recursive and canonical splicing sites, we generated heatmaps for relative the log2 fold change of R-values, where the canonical splicing sites show the ratio of R-values from introns with RS events and introns without RS events, and the recursive splicing sites show the ratio of R-values between recursive and canonical splicing site of RS introns (Figure 7B, S5). At the regions flanking the first canonical 5’SS and 3’SS, which contain the vital features for RS prediction, we observe several known splicing factors among the top hits, including RBM11, hnRNPC, PCBP1/2, SFPQ and SF3B6, with potential differential binding activities evidenced as opposite enrichment trends at the canonical and recursive SS (Figure 7B, S5). For example, RBM11 is depleted at the canonical 5’SS but enriched near the recursive 5’SS and shows negative preference at the recursive 3’SS and positive preference at the canonical 3’SS (Figure 7B, S5). We speculate that potential competition for the same splicing factors might distinguish canonical and recursive SS. These candidates and other RBPs identified in this analysis constitute a computational prediction that can be tested in future experimental studies.

**Figure 7.**
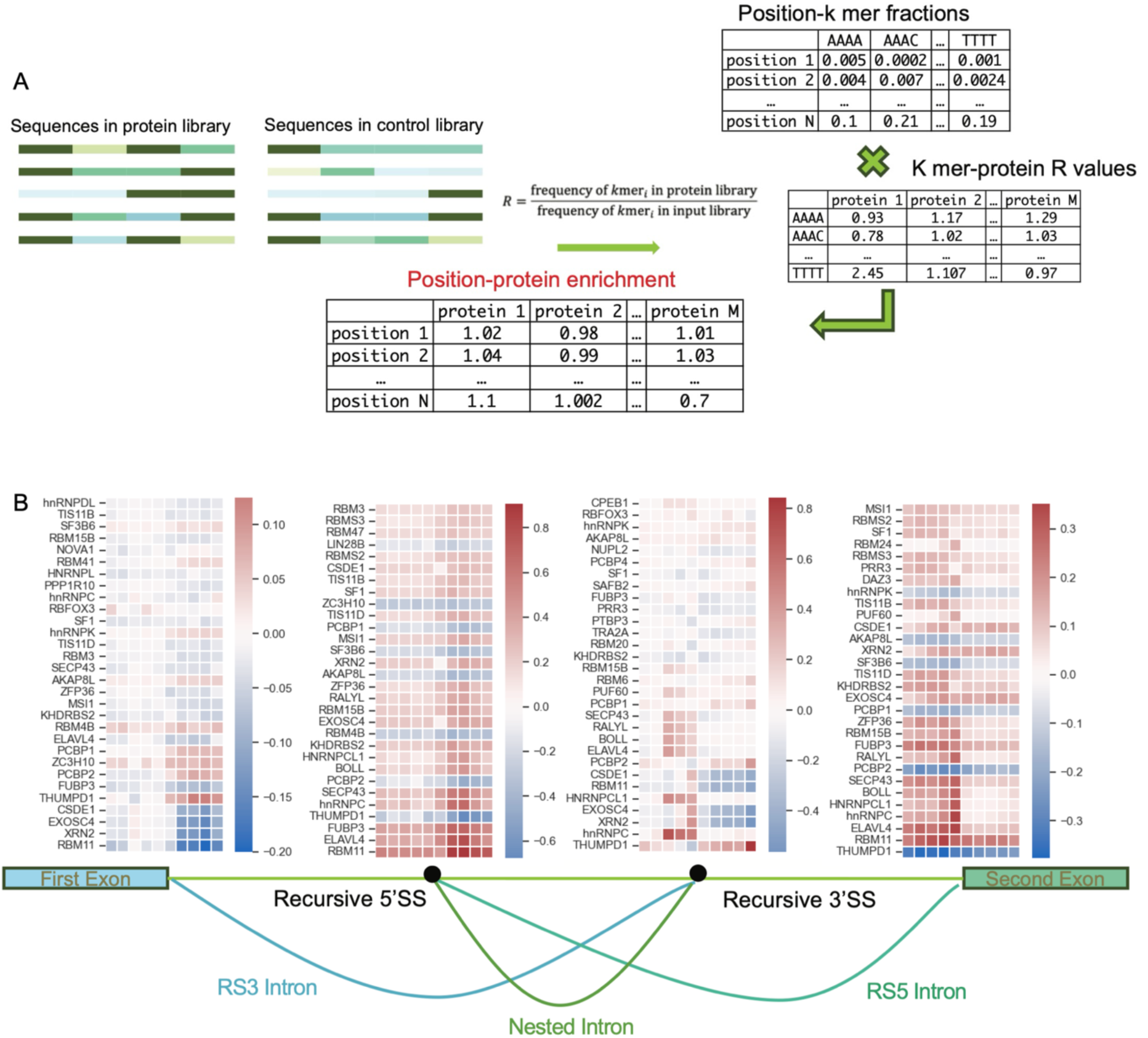
In vitro protein binding assay uncovers potential trans acting RBPs regulating recursive splicing. A. Evaluation of protein enrichment by matrix multiplication. R values of RBNS experiments available from ENCODE are used to make a k-mer by protein matrix. K-mer counts from nascent RNA-Seq sequences are used to make a position by k-mer matrix. Matrix multiplication produces position by R-value matrix, showing RBP binding potential of different positions of introns with recursive splicing. B. Heatmap colored by relative protein enrichment. The top30 log2FC values of protein enrichments based on 5mer RBNS are shown. (At the canonical 5’SS and canonical 3’SS, the enrichment=enrichment in RS introns/enrichment in no-RS introns; at the recursive 5’SS and recursive 3’SS, enrichment=enrichment at recursive sites in RS introns/enrichment at canonical sites in RS introns; 1 cube=15bp. Proteins are ranked by the highest absolute L2FC in each protein.)

## Discussion

Recursive splicing appears to be ubiquitous in human RNA processing, begging the question: how is this information encoded in the genome? By applying sequence analysis algorithms, such as Latent Dirichlet Allocation, to the nascent RNA-Seq data, we have identified sequence features, one enriched in CG dinucleotides located in the flanking region of the canonical 5’SS of the first introns, and one with purine dilution in the polypyrimidine tract region of the canonical 3’SS of the first introns, associated with RS. Genes with these features show higher rates of recursive splicing not only in the first intron but also in downstream introns, making these features a predictor of recursive splicing throughout the transcription unit. The recursive splicing classifier we develop has > 85% accuracy in the first introns and >75% accuracy in downstream introns and relies on features which provide potential mechanistic insight into the larger question of splice site selection in human cells.

### Mixture models can identify features from RNA sequences

Latent Dirichlet allocation is a generative probabilistic model in natural language processing that summarizes topics from a collection of documents (Blei et al., 2003). These approaches have shown great promise in sequence analysis and have been used previously for population structure inference and even applications in RNA biology (Blei et al., 2003; Dey et al., 2017; Pritchard et al., 2000; B. Wang & Mount, 2024). Here, we applied the LDA mixture model to k-mer compositions of recursive splicing sequences in two ways: searching for features of grouped sequences at different locations relative to the splicing sites and searching for features in single sequences. With both methods, we observed that the region flanking the canonical 5’SS of recursively spliced introns at the beginning of genes is enriched in a collection of k-mers, where all the top-ranked ones contain CG-dinucleotides. Also, the region flanking the canonical 3’SS of the recursively spliced introns shows lower enrichment of k-mers that contain CTs, which is associated with a strong polypyrimidine tract but increased AT-rich k-mers. When coupled with a random forest classifier, the motifs discovered by mixture models provide better predictions of recursive splicing than a simple GC content measurement, suggesting that LDA can identify more detailed and subtle sequence features.

However, the topics/features calculated by LDA are represented as mixtures composed of the probabilities of observing each possible k-mer in each topic or feature, which are not suitable for making sequence logos. Whether ‘topics’ or ‘logos’ are better at capturing biological reality in this use case is unknown. It is possible that RS is generated by a class of RBPs all of which contribute to the outcome and bind generally similar sequences as opposed to a single RBP determining the outcome. For example, one RBP we identified in this analysis, SFPQ, was shown to be involved in long intron splicing, which would support our analysis (Stagsted et al., 2021). Yet, it might just be one of many RBPs playing a role. In this case, one might not identify a logo/position weight matrix which corresponds to a specific RBP site. It is also possible that the CG preference and a different polypyrimidine tract correspond to physical properties of the DNA/RNA which are determining splicing outcomes. There is a long history of interrogating GC content to understand downstream consequences of gene expression including splicing (Barutcu et al., 2022; Lemaire et al., 2019; Tammer et al., 2022), RNA localization (Mordstein et al., 2020; Tammer et al., 2022) and export (Zuckerman et al., 2020) as some recent examples. These two models – RBP binding or thermodynamic/global effects – are of course not mutually exclusive. We also noticed that many RBPs show opposite binding activities between the canonical and recursive SS. For example, both PCBP1 and PCBP2 show depleted enrichment at the canonical 3’SS, but increased binding activities at the recursive 3’SS, which is consistent with the feature distribution inferred by the mixture model that recursively spliced introns have fewer C-rich k-mers flanking the canonical 3’SS. Both PCBP1 and PCBP2 are involved in alternative splicing regulations (Ji et al., 2018), and the opposite enrichment patterns between different 3’SS could explain splice site usage in the RS model.

### Features of recursive splicing

We observed that a large fraction of recursively spliced introns in the human genome are first introns, which are typically more than twice as long as downstream introns. Therefore, recursive splicing may contribute to splicing fidelity by allowing a very long intron to be removed through small pieces (Duff et al., 2015; Gehring & Roignant, 2021; Hatton et al., 1998; Pai et al., 2018; Wan et al., 2021). Several studies have shown that RS proceeds via exon definition through defining a transient RS exon, which is skipped thereafter (Joseph et al., 2018; Pai et al., 2018; Sibley et al., 2015). Yet, splicing can also occur independently of exon definition (Zeng et al., 2022). Interestingly, we observed differences in GC content in the intron sequences upstream and downstream of potential recursive sites, which are similar to the exon and intron definition-related gene architecture identified by a previous study (Tammer et al., 2022). At recursive 3’SS, we observe higher GC content downstream than in the upstream sequences, which is consistent with the GC content architecture of exon definition. One explanation is that the downstream sequence is recognized as an RS exon, mimicking exon definition before the correct downstream exon is transcribed. While at recursive 5’SS, the difference is smaller, and both upstream and downstream intron subsequences show higher GC content than the upstream subsequence of recursive 3’SS, which looks like the architecture of intron definition that allows a smaller RS intron to be spliced first. Our observations suggest that both exon and intron definitions have the potential to initiate recursive splicing site usage.

Our results imply that a preference for recursive splicing sites is established at the beginning of transcription through a CG-rich motif and a potentially different polypyrimidine tract. This conclusion is based on two observations. First, Fisher’s Exact test indicates that the likelihood of observing recursively spliced downstream introns in a gene with a recursively spliced first intron is significantly higher than in a gene with a constitutively spliced first intron. Second, when grouping genes based on splicing patterns in the downstream introns, sequences flanking the canonical 3’SS of the first introns from genes with downstream recursive splicing events are more enriched in the AT-rich motif, the same motif of recursive splicing in the first introns, even when the splicing patterns of the first introns are unknown. Additionally, sequences flanking the 5’SS of the first introns with RS evidence are more enriched in a CG-rich motif than canonical introns. Taken together, we favor a biochemical model where RBPs are recruited by CG-rich sequences and AT-rich polypyrimidine tracts and then regulate downstream splicing activity for the whole transcript. Indeed, our analysis of ENCODE RBNS data provides several compelling predictions to be tested experimentally.

Interestingly, whole-genome bisulfite sequencing reveals that the enrichment of the CG-rich motif of recursively spliced first introns is coupled with a low CpG methylation status in the first exons compared with regularly spliced first introns (Dunham et al., 2012; Luo et al., 2020). We envision two possible explanations for this result. Previous studies have revealed that DNA methylation can affect RNA splicing through multiple mechanisms, including modulation of RNA polymerase II elongation rate and recruitment of splicing factors via methyl-binding proteins and histone modifications (Agirre et al., 2021; Anastasiadi et al., 2018). Thus, potential recursive splicing regulators could be recruited through the same mechanism. On the other hand, methylated CpG has reduced chemical stability and eventually mutates (Pértille et al., 2019; Tomkova et al., 2024), leading to sequence drift over time. This fact suggests an alternative explanation: selection for preservation of the CG-rich motif favors a low methylation rate. We favor this latter explanation based both on established splicing biochemistry and also the predicted differential binding of RBPs in the CG-rich region. Overall, we propose a mechanism in which early events in RNA synthesis and processing influence the prevalence of recursive splicing for the entire transcript, potentially activating an alternative version of the spliceosome.

### Limitations of the study

The nascent RNA-Seq and lariat sequencing provide a solution to splicing intermediate detection, as well as challenges to the investigation of recursive splicing mechanisms. Due to the various speeds of splicing and the possibility that splicing can happen within seconds, many sequencing events cannot be captured. Thus, the background intron sequences that were labeled as basic introns contain an undetermined number of false negatives. Our study is exclusively based on nascent RNA-Seq of cultured human cells, and the recursive splicing pattern may differ in different tissues and cell types. Though our results reveal candidate RBPs that have a preference for sequences associated with recursive splicing, they may not play any role in the recursive splicing regulations, while other RBPs that are absent from the RBNS catalog can interact with recursive SS through the same motifs.

## Methods

### Identification of potential recursive splicing sites

First, to avoid mixing recursive splicing with alternative splicing site usage, we select constitutive introns from the human genome by discarding those that appear in annotated alternative isoforms from the RefSeq GTF annotations and those that could potentially overlap with any other introns in the human genome using bedtools intersect (Goldfarb et al., 2025; Quinlan & Hall, 2010).

Next, we search for potential recursive splicing events. From STAR splice junction output files, we take all splicing junctions supported by at least 3 unique reads (Dobin et al., 2013). We compare the coordinates of the splicing junctions with the list of constitutive introns and select 4 groups for further analysis: (1) basic introns where both splice sites align with corresponding annotated introns, (2) RS5 introns where the 5’SS is downstream of the annotated 5’SS and the 3’SS is the annotated 3’SS of the constitutive intron, (3) RS3 introns where the 3’SS is upstream of the annotated 3’SS and the 5’SS is the annotated 5’SS of the constitutive intron it overlaps with, (4) nested introns where the 5’SS is downstream of the annotated 5’SS and the 3’SS is upstream of the annotated 3’SS of the overlapping constitutive intron. We use basic introns at least 75bp long and the 3 types of recursive introns where their corresponding constitutive introns are at least 150bp long with recursive splice sites at least 75bp away from the annotated splice sites. We did this for both nascent RNA-Seq and regular RNA-Seq and use the splicing junctions that are uniquely found in nascent RNA-Seq as potential recursive splicing events.

### Fitting mixture models for sequence analysis

We fit mixture models using a sequence feature analysis Python library based on sci-kit learn (Harris et al., 2020; Hunter, 2007; Pedregosa et al., 2011; B. Wang & Mount, 2024). First, we explore features associated with positions relative to the splice sites. We take sequences from 30bp upstream to 50bp downstream of the canonical and recursive 5’SS and label them based on the intron type. For each intron and 5’SS subtype, we count the tetramers at each position with a window of 8bp, which helps us create a matrix where each column is a label containing information on position, intron type, 5’SS type, and each row is a tetramer. Mixture models are fitted using the matrices that calculate the topic memberships in the data. Next, we test if the feature discovered by the previous method is distinctively found in recursive splicing or is driven by a special subset or outliers. We extract sequences from the 50 bp upstream to the 50 bp downstream of each 5’SS and count tetramers within each sequence. For each intron subtype, we randomly select 100 sequences and combine the tetramer counts as a single sample to avoid overfitting and low interpretability of the topics. We determined the number of topics based on the performance of the LDA model in distinguishing recursive and canonical SS.

We calculate the driving k-mers of each topic by calculating the distinctiveness of each tetramer using Kullback-Leibler divergence (Dey et al., 2017; B. Wang & Mount, 2024).

### DNA CpG methylation calculation

We used A549 cell line WGBS data provided by ENCODE to calculate average exon CpG methylation (Dunham et al., 2012; Luo et al., 2020). For each first exon from our dataset, we collect all the CpGs from the sequence and calculated the average methylation percentage.

### Random forest classifiers for recursive splicing predictions

To prepare training data, we fit mixture models to measure topic memberships in regions upstream and downstream of the canonical 5’SS and 3’SS for both first and downstream introns. We take splicing junctions with at least 3 unique supporting reads from transcripts with TPM>=1 to build the training and test sets.

We fit random forest classifiers using Sci-Kit learn (Pedregosa et al., 2011; Virtanen et al., 2020). We randomly select 1/3 of the data as the test set. For each of the classifiers we tested for the first introns, we used hyperparameters of n_estimators, min_samples_split, min_samples_leaf, max_depth as [100,10,5,3] and we used [100,10,5,2] for downstream intron tests. ROC curve and precision-recall curve are calculated with scipy and Sci-Kit learn packages.

### Recursive splicing validation by LSV-seq

LSV-seq primers were designed as previously described (Yang et al., 2025), targeting exons downstream of control or candidate introns, and ordered as a single oligo pool (oPools, Integrated DNA Technologies). Nuclei from HBEC3-KT cells were isolated using the REAP protocol (Suzuki et al., 2010) and RNA was extracted using the Direct-zol RNA Purification Kit (Zymo Research, D2052). LSV-seq libraries were prepared from 5 ug of input RNA as previously described (Yang et al., 2025), with the following minor modifications. The final PCR amplification step was performed with NEBNext Ultra II Q5 Master Mix (NEB, M0544). After an initial 0.9X AMPure XP bead cleanup, libraries were run on a 2% EX E-Gel (Invitrogen, G401002) and further purified by extraction of the 200-600bp range using the Zymoclean Gel DNA Recovery Kit (D4008, Zymo Research). Finally, libraries were pooled and sequenced in 150-cycle single-end format on an Illumina NextSeq 550.

LSV-seq data was processed as previously described (Yang et al., 2025). Briefly, reads were first trimmed with BBDuk of BBMap (sourceforge.net/projects/bbmap/) and UMIs were extracted from each read using umi-tools (Smith et al., 2017). Reads were then aligned to the hg19 genome with STAR (Dobin et al., 2013) and the resulting BAM files were deduplicated with UMICollapse (Liu, 2019) and a custom python script.

### Identification of potential trans-acting factors of recursive splicing

To search for RBPs that are associated with the recursive splicing sequences, we downloaded the RNA Bind-N-Seq k-mer enrichment data from ENCODE (Dominguez et al., 2018a; Dunham et al., 2012; Luo et al., 2020; Van Nostrand et al., 2017). We pick the concentration with the highest enrichment value for each protein and create a protein-6-mer matrix. In both recursively-spliced introns and constitutive introns, we also count the 6-mers of the same length at every position relative to the splice site, generating a sequence-6-mer matrix. We multiply the matrices to get enrichment of RBPs at different positions of the sequences and calculate the ratio between recursive splicing and constitutive introns.

## Acknowledgment

We thank members of the Larson and Mount laboratories for helpful discussions. We want to specifically thank Benjamin Donovan, Yihan Wan, and Jee Min Kim for insightful discussions on splicing mechanisms. We thank members of the Chris Burge laboratory for valuable feedback. We acknowledge the use of GitHub Copilot for coding assistance. This work was supported by Biowulf, the High-Performance Computing Group, and the Center for Information Technology. United States Government disclaimer: this work was supported by the NIH Intramural Research Program through award 1ZIABC011383. The contributions of the NIH author(s) were made as part of their official duties as NIH federal employees, are in compliance with agency policy requirements, and are considered Works of the United States Government. However, the findings and conclusions presented in this paper are those of the author(s) and do not necessarily reflect the views of the NIH or the U.S. Department of Health and Human Services.

## Supplementary Figures

**Supplementary Figure 1.**
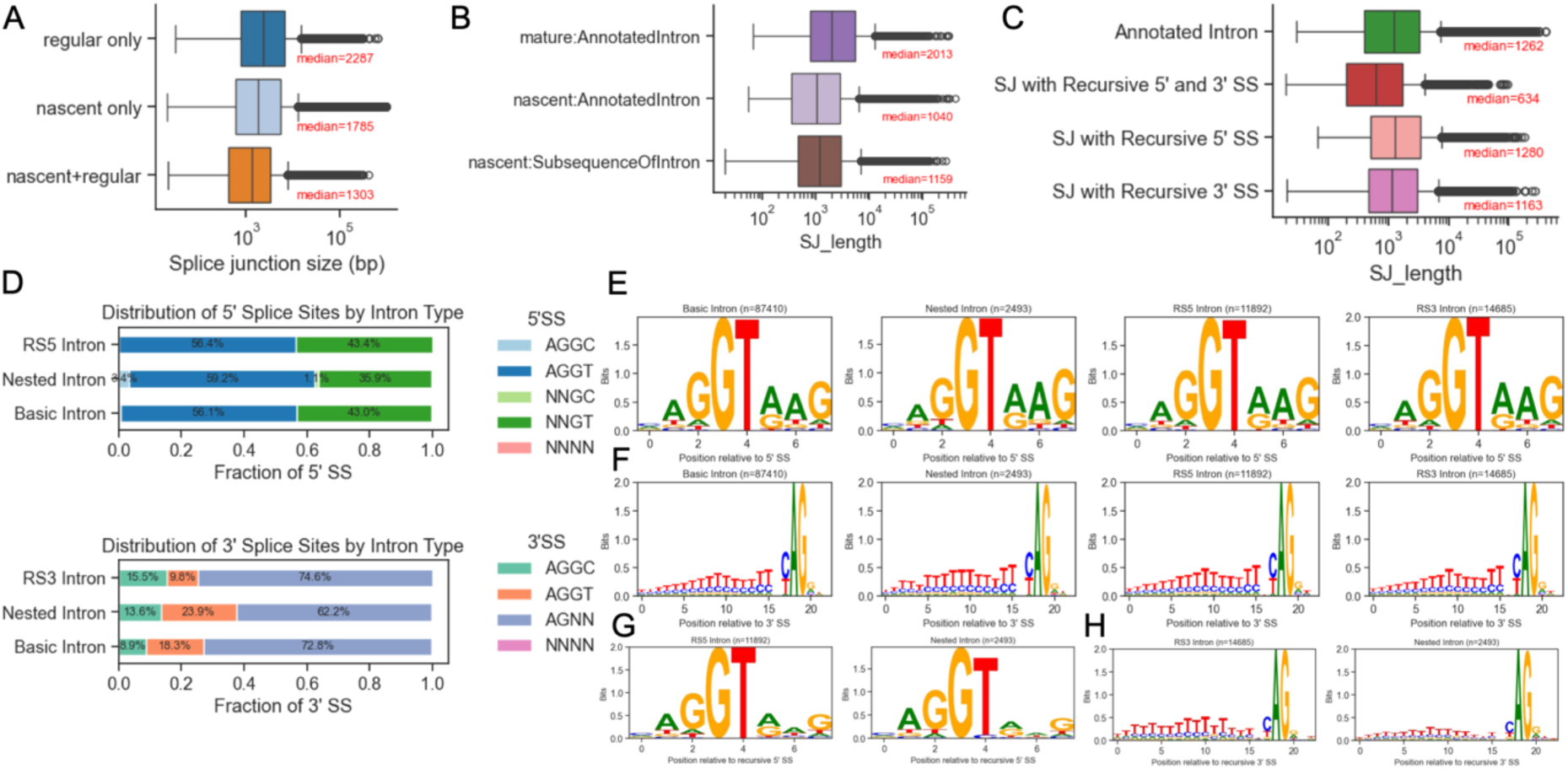
Nascent RNA-Seq revealed 4 types of introns. A. Distribution of unique splice junctions in either nascent or regular RNA-Seq and those ubiquitously used in both data. B. Distribution of lengths of splice junctions uniquely expressed in either nascent or regular RNA-Seq. C. Distribution of splice junction lengths in 4 types of introns. D. Fractions of 5’SS tetramers and 3’SS tetramers at recursive splice sites. E. Sequence logo of annotated 5’SS of 4 types of introns. F. Sequence logo of annotated 3’SS of 4 types of introns. G. Sequence logo of recursive 5’SS of 2 types of introns. H. Sequence logo of recursive 3’SS of 2 types of introns.

**Supplementary Figure 2.**
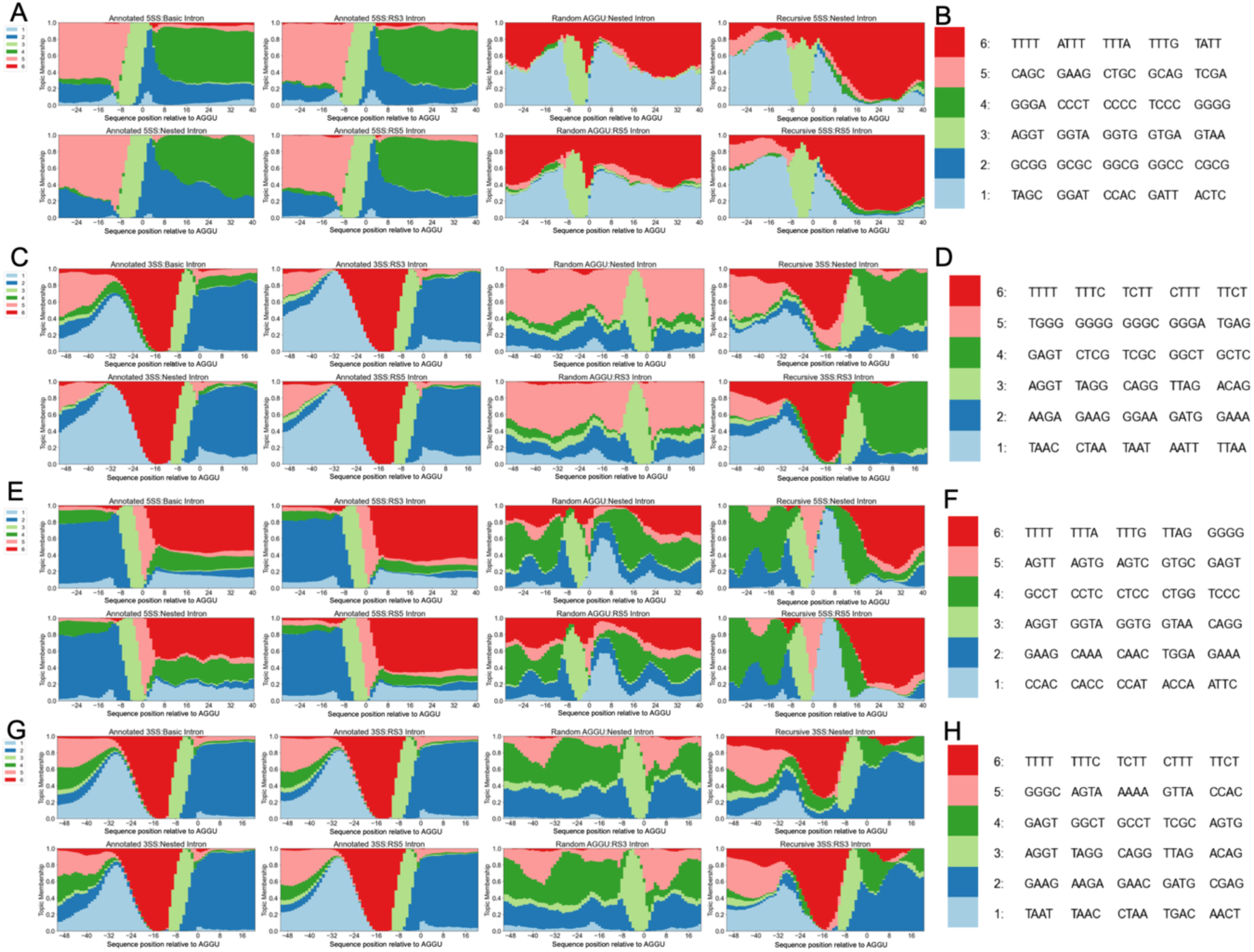
Mixture models uncovered potential cis-acting factors of recursive splicing. A. Structure plot of topic/cluster distribution of regions from 30bp upstream to downstream 50bp of canonical and recursive 5’SS in the first introns. Random AGGU sites were also shown. B. Top ranked k-mers in 5’SS model in A. C. Structure plot of topic/cluster distribution of regions from 50bp upstream to 30bp downstream of 3’SS in the first introns. D. Top ranked k-mers in 3’SS model in C E. Structure plot of topic/cluster distribution of regions from 30bp upstream to downstream 50bp of canonical and recursive 5’SS in the downstream introns. Random AGGU sites were also shown. F. Top ranked k-mers in 5’SS model in E. G. Structure plot of topic/cluster distribution of regions from 50bp upstream to 30bp downstream of 3’SS in the downstream introns. H. Top ranked k-mers in 3’SS model in G.

**Supplementary Figure 3.**
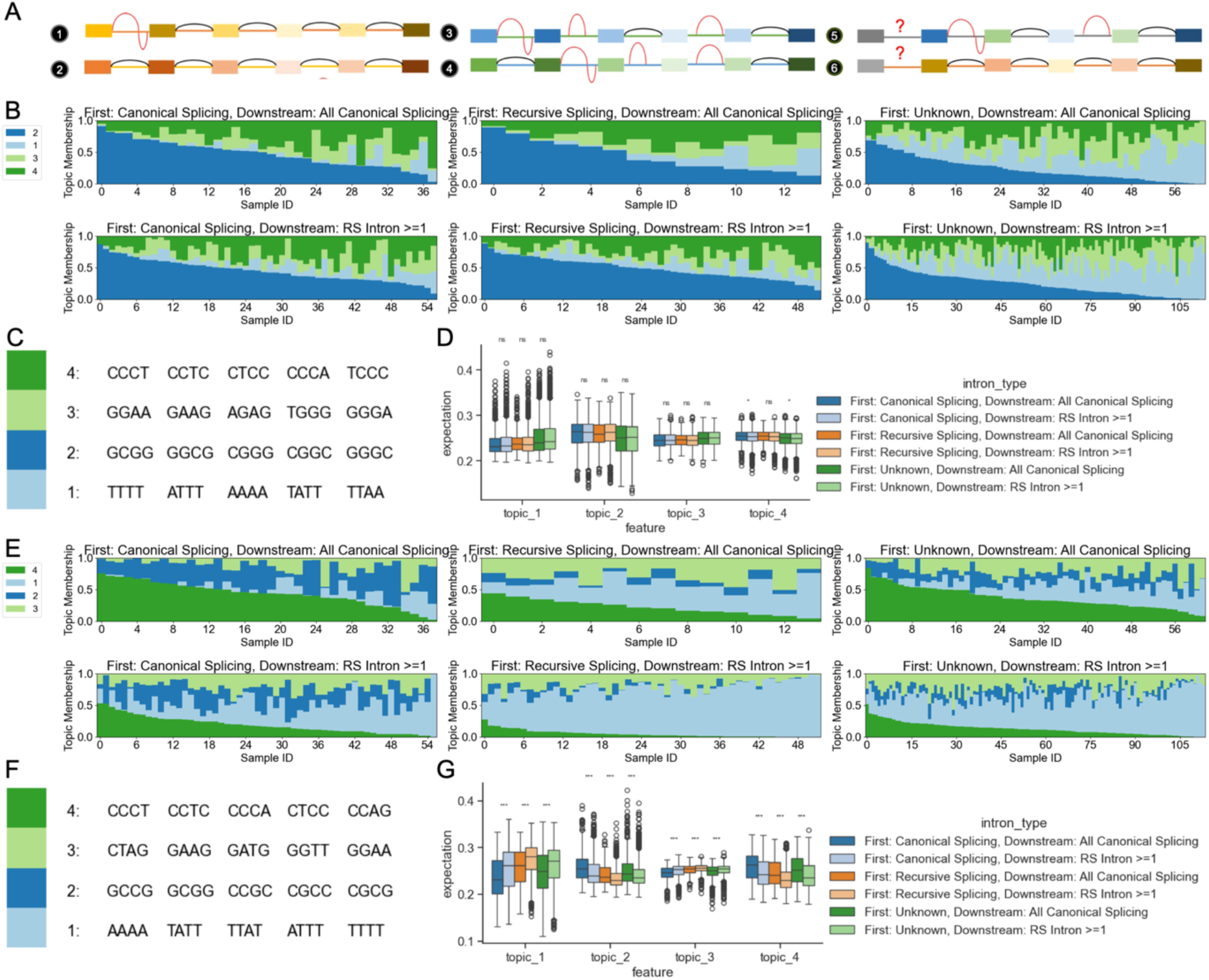
Sequences of the annotated 5’SS and 3’SS of the first intron indicate recursive splicing site usage. A. Diagram of 6 types of transcription units based on the splicing status in the first and downstream introns. B. Structure plot of topic/cluster distribution of regions from 50bp upstream to 50bp downstream of the canonical 5’SS in the first introns. C. Top ranked k-mers in 5’SS model in B. D. Distribution of single sequence topic expectation values calculated with model B. E. Structure plot of topic/cluster distribution of regions from 50bp upstream to 50bp downstream of the canonical 3’SS in the first introns. F. Top ranked k-mers in 3’SS model in E. G. Distribution of single sequence topic expectation values calculated with model E.

**Supplementary Figure 4.**
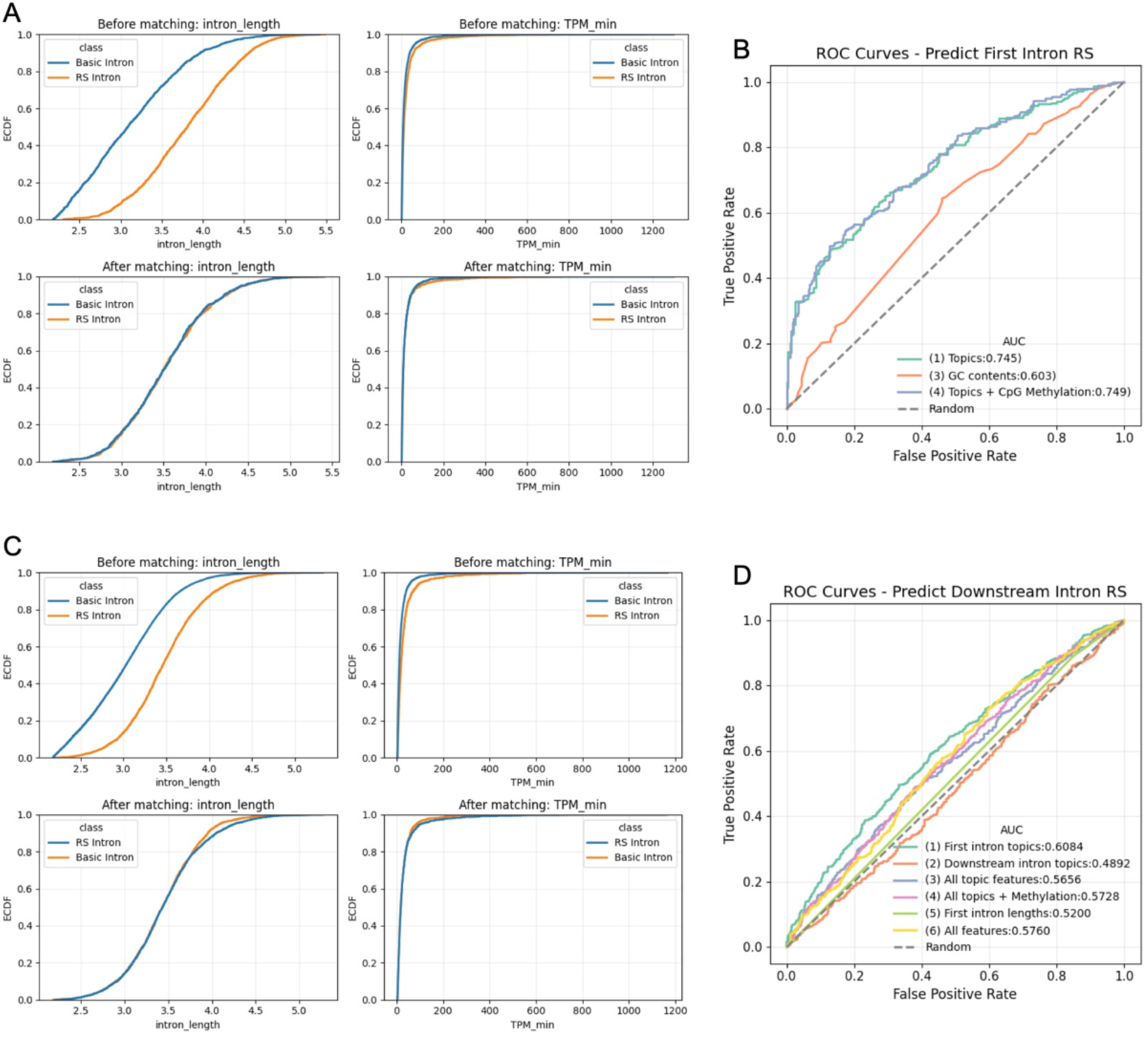
Computational modeling of sequence features aids in the accurate prediction of recursive splicing. A. Cumulative distribution of intron length and TPM of basic and RS introns (first introns of genes) before and after adjusting matching sets. To evaluate the the performance without the impact of intron length or gene expression, we select subsets of basic and RS first introns with similar length and TPM distribution. B. ROC curve of predicting RS in first introns. Mixture model topics and DNA methylation can help predict RS with accuracy of 74.9% C. Cumulative distribution of intron length and TPM of basic and RS introns (downstream introns of genes) before and after adjusting matching sets. D. ROC curve of predicting RS in downstream introns. Sequence features in the corresponding first introns help predict RS in downstream introns of the same genes with an accuracy of 60.8%.

**Supplementary Figure 5.**
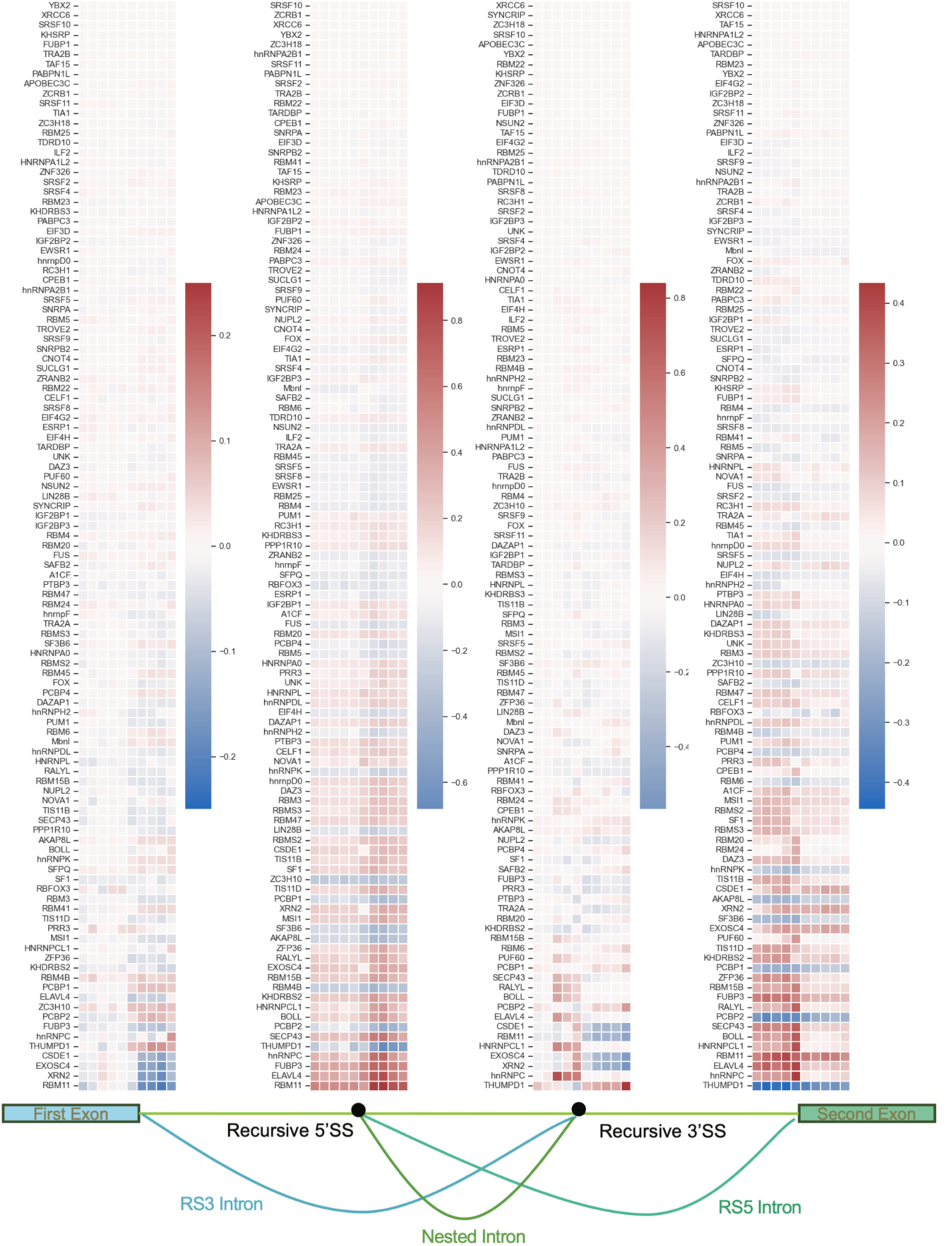
In vitro protein binding assay uncovers potential trans acting RBPs regulating recursive splicing. Full heatmaps of the L2FC of protein-binding enrichment scores in F7B.

